# Cross-Platform Evaluation of a Tumor-Budding-Associated Gene Signature in Colorectal Cancer

**DOI:** 10.64898/2026.10.09.757976

**Authors:** Pritom Sarkar, Prapthi Sarkar

**Author notes:** **Emails:** (Corresponding Author) (Second Author).

## Abstract

Spatial transcriptomics has identified *AIF1, LIF, MRC1*, and *TNC* as enriched in tumor-budding (TB) regions of colorectal cancer. We investigated whether this TB-associated gene module could be detected in bulk transcriptomic data. Pairwise gene co-occurrence was analyzed in bulk RNA-seq (TCGA-COADREAD, n=547) and single-cell RNA-seq (GSE178341, n=26,057 cells) using Fisher’s exact tests with Benjamini-Hochberg correction. In bulk RNA-seq, pairwise fold-enrichments were weak (1.16–2.72x), with *AIF1–LIF* showing no significant association (FE=1.16x, FDR=0.31). In single-cell data, the same pairs showed 31–184.5x (all FDR < 8×10⁻³⁸). Total UMI counts provided no evidence of doublet artifacts explaining these co-detection patterns. Exploratory enhancer–promoter interaction analysis in HCT116 cells further suggested these genes are controlled differently. *GOT1* and *MDH1*, two malate–aspartate shuttle genes, showed lower expression in late-stage CRC (FDR ≤ 0.010). A bulk-derived risk score showed a modest association (Cox HR=1.21, p=0.021). Overall, TB-associated co-expression was more clearly detected at single-cell than bulk resolution, highlighting the importance of transcriptomic resolution when studying tumor-microenvironment-associated gene signatures.

## Introduction

Colorectal cancer is a major cause of cancer-related deaths worldwide.^1,2^ A feature of advanced CRC is tumor budding, in which single cancer cells or small clusters detach from the invading edge of the tumor, and is associated with poor prognosis in CRC. Understanding the molecular features associated with tumor budding may therefore provide important insights into CRC progression.^3,4^ Tumor budding signals a poor outcome in colorectal cancer, but the molecular features associated with tumor budding remain incompletely understood.^5^ Recent spatial and single-cell transcriptomic studies have provided insights into the cellular heterogeneity of the CRC tumor microenvironment.^6,7^ However, it remains unclear whether gene-expression patterns associated with spatially restricted tumor budding regions can be detected in bulk RNA-seq.

Recently, an AI-based virtual spatial transcriptomics tool named Phoenix identified a specific four-gene signature(*AIF1, LIF, MRC1*, and *TNC*) that is significantly enriched in these tumor budding (TB) zones, forming a reactive tumor microenvironment (TME) that is completely different from the normal, stable programs of nearby tumor regions.^8^ This discovery raises an important translational question: can these TB-associated co-expression patterns be detected using bulk RNA sequencing?

Bulk RNA-seq measures the average gene expression across millions of cells of diverse types within a tissue sample, including tumor, immune, stromal, and endothelial cells. Although this approach is useful for studying overall gene expression, it may be less sensitive to signals from rare or spatially restricted cell populations. This averaging process can easily hide signals that come from tightly localized cell groups. The statistical consequence is that gene-gene correlations observed at the whole-tissue (bulk) level may simply reflect variation in the cell-type mixture, rather than true co-regulation within individual cells.^9,10^ When the specific cells of interest make up only a tiny fraction of the total tissue, as is the case for TB-associated immune populations, these bulk-level signals are expected to be severely weakened compared to single-cell measurements.

In this study, we systematically test our hypothesis. First, we calculate the co-occurrence of the TB-module genes in TCGA-COADREAD bulk RNA-seq data (n=547) using pairwise Boolean co-detection analysis backed by statistical testing (Fisher’s exact test with BH-FDR correction). Second, in HCT116 colorectal cancer cells using UniversalEPI, we map the predicted enhancer–promoter interaction landscape of these genes, providing crucial chromatin-level regulatory context.^11^ Third, we examined the single-cell co-expression structure of the TB module in an aggressive-subtype CRC single-cell RNA-seq dataset (n=26,057 cells),^12^ while applying strict controls to eliminate doublet artifacts.^13^ Fourth, we examine how MAS pathway and glutamine metabolism genes change across CRC clinical stages.^14^ Together, our results demonstrate that bulk RNA-seq completely fails to catch pairwise co-expression patterns that are robustly visible at single-cell resolution, providing an example of how tissue-level averaging masks cell-type-specific biology in colorectal cancer.

## Result

### Promoter region extraction

The 2,500 bp promoter intervals for all eight candidate genes were mapped using the Ensembl GRCh38 annotation. We mapped the exact genomic regions relative to the Transcription Start Sites (TSS) to establish a standard baseline for downstream motif and regulatory analysis. (Supplementary Table 1).

### Characterization of the Predicted Enhancer-Promoter Interaction (EPI) Landscape in HCT116 Cells

To understand how our candidate genes (*AIF1, TNC, LIF, MRC1, GOT1, MDH1, GOT2, MDH2*) are regulated, we explored their predicted Enhancer-Promoter Interaction landscapes specifically in the HCT116 colorectal cancer cell line. Glutamine metabolism genes (*GLS*, *GLUD1*, and *GLUL*) were excluded because they did not show significant differential expression between clinical stages (FDR > 0.05; see Table 4). We mapped the location of all predicted interaction anchors and calculated their statistical significance using a Z-score threshold (Z > 1.645). Table 1 shows that many genes exhibit a highly active regulatory landscape. *GOT1, TNC, MDH2*, and *LIF* displayed the strongest interactions. *GOT2, MDH1*, and *MRC1* showed fewer active interactions, and interestingly, *AIF1* had no predicted anchors because its entire promoter bin was absent from the HCT116 EPI map (Supplementary Figure S1).

**Table 1.**
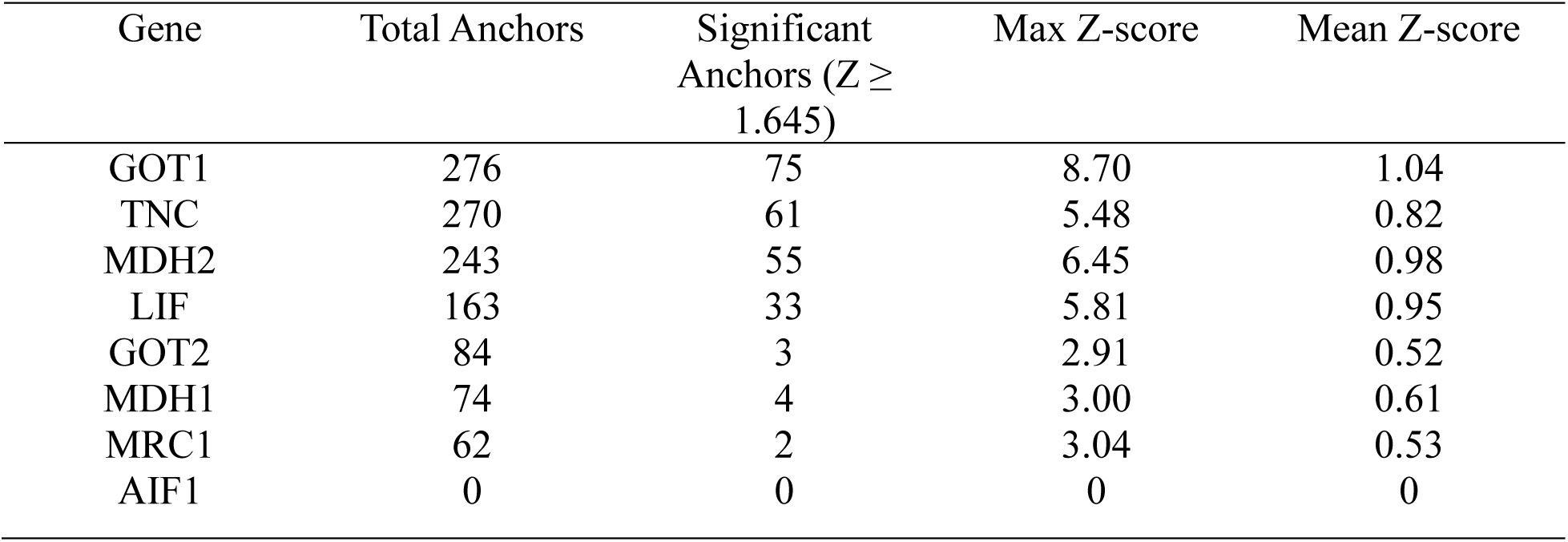
Promoter coordinates (GRCh38)

Since this characterization is based on a single cell line (HCT116; n = 1), these results are presented as exploratory findings rather than definitive statistical claims.

### Bulk RNA-seq: activation rates and pairwise co-occurrence

We analyzed bulk RNA-seq data from the TCGA-COADREAD cohort (n=547; Early Stage I–II, n=320; Late Stage III–IV, n=227). For each gene, samples with expression values above the 75th percentile were classified as HIGH expression and the remaining samples classified as LOW expression. The 75th percentile thresholds are shown in Table 2.

**Table 2.** Threshold.

|  |  |
| --- | --- |
| AIF1 | 253.88 |
| TNC | 2984.37 |
| LIF | 1043.78 |
| MRC1 | 332.55 |

Using this threshold, approximately 22–27% of samples showed HIGH expression for each of the four genes across both tumor stages, as shown in Figure 1a, with no clear stage-dependent separation.

**Figure 1a.**
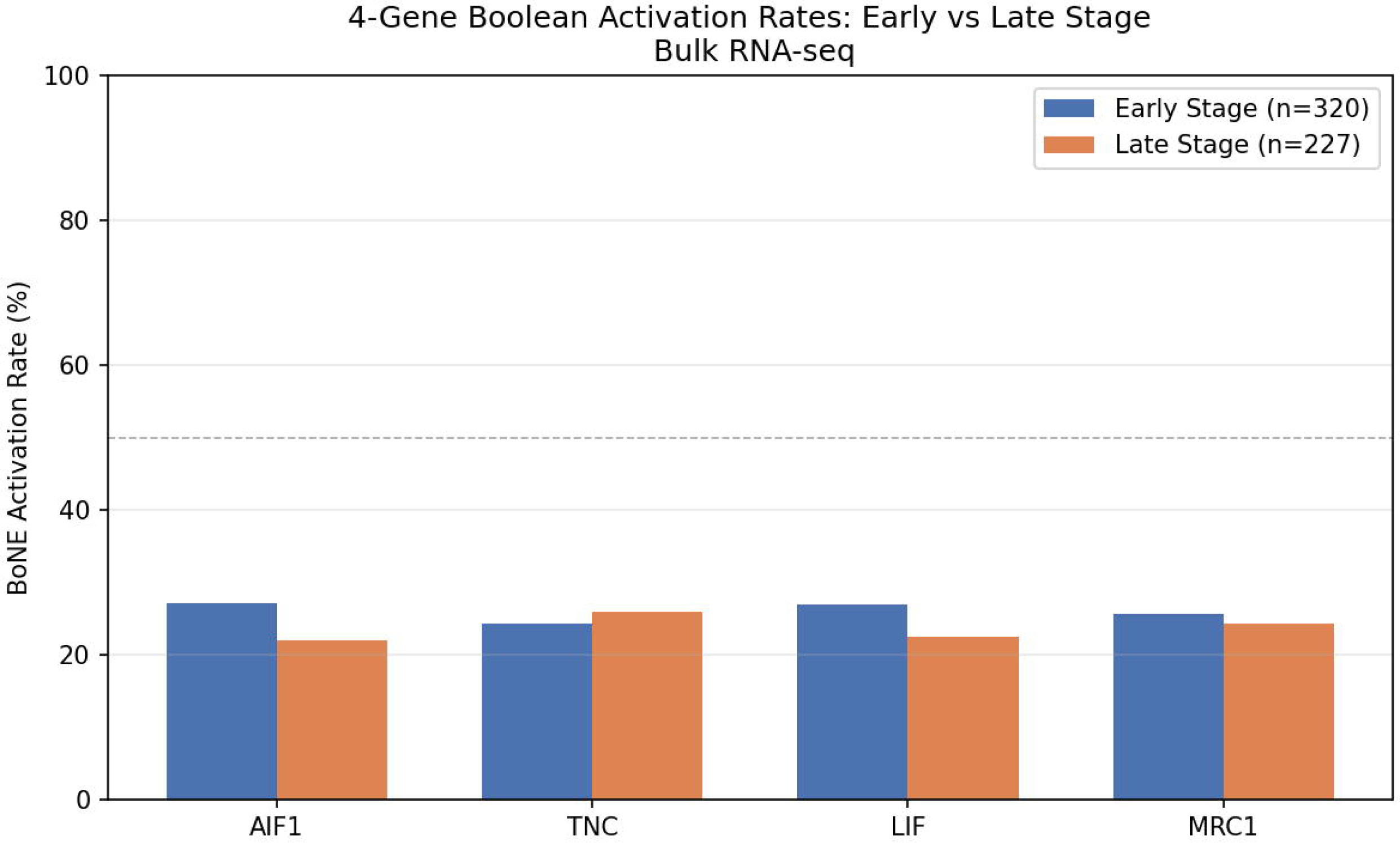
Tumor-budding module activation rates across CRC disease stages.

To quantify pairwise gene co-occurrence in bulk tissue, we used Fisher’s exact test to assess whether all 12 directional gene pairs were co-expressed in late-stage TCGA-COADREAD samples. Multiple testing was corrected using the Benjamini–Hochberg false discovery rate (FDR) method (Table 3). The pairwise fold enrichment ranged from 1.16 to 2.72. The strongest co-occurrence was observed between *AIF1* and *MRC1* (fold enrichment = 2.72, FDR = 1.7 × 10⁻¹²), suggesting frequent co-expression of these two macrophage-related genes. Notably, *AIF1* and LIF did not show a statistically significant association (fold enrichment = 1.16, FDR = 0.31), despite being a known macrophage-secreted cytokine. *TNC* showed moderate but significant co-expression (fold enrichment = 1.66–2.17; all FDR < 0.004; Figure 1b).

**Table 3.** Bulk pairwise Fisher’s and FDR.

| Primary Gene (IF) | Target Gene (THEN) | Precision (%) | Fold Enrichment | Odds Ratio (OR) | p-value | FDR |
| --- | --- | --- | --- | --- | --- | --- |
| AIF1 | MRC1 | 66.0 | 2.72 | 13.68 | $2.89 \times 10^{-13}$ | $1.73 \times 10^{-12}$ |
| MRC1 | AIF1 | 60.0 | 2.72 | 13.68 | $2.89 \times 10^{-13}$ | $1.73 \times 10^{-12}$ |
| MRC1 | TNC | 56.4 | 2.17 | 6.64 | $1.98 \times 10^{-8}$ | $5.95 \times 10^{-8}$ |
| TNC | MRC1 | 52.5 | 2.17 | 6.64 | $1.98 \times 10^{-8}$ | $5.95 \times 10^{-8}$ |
| AIF1 | TNC | 52.0 | 2.00 | 4.73 | $5.97 \times 10^{-6}$ | $1.19 \times 10^{-5}$ |
| TNC | AIF1 | 44.1 | 2.00 | 4.73 | $5.97 \times 10^{-6}$ | $1.19 \times 10^{-5}$ |
| LIF | TNC | 43.1 | 1.66 | 2.85 | $1.86 \times 10^{-3}$ | $2.78 \times 10^{-3}$ |
| TNC | LIF | 37.3 | 1.66 | 2.85 | $1.86 \times 10^{-3}$ | $2.78 \times 10^{-3}$ |
| LIF | MRC1 | 35.3 | 1.46 | 2.05 | $3.05 \times 10^{-2}$ | $3.67 \times 10^{-2}$ |
| MRC1 | LIF | 32.7 | 1.46 | 2.05 | $3.05 \times 10^{-2}$ | $3.67 \times 10^{-2}$ |
| AIF1 | LIF | 26.0 | 1.16 | 1.29 | 0.31 | 0.31<br>(NS) |
| LIF | AIF1 | 25.5 | 1.16 | 1.29 | 0.31 | 0.31<br>(NS) |

**Figure 1b.**
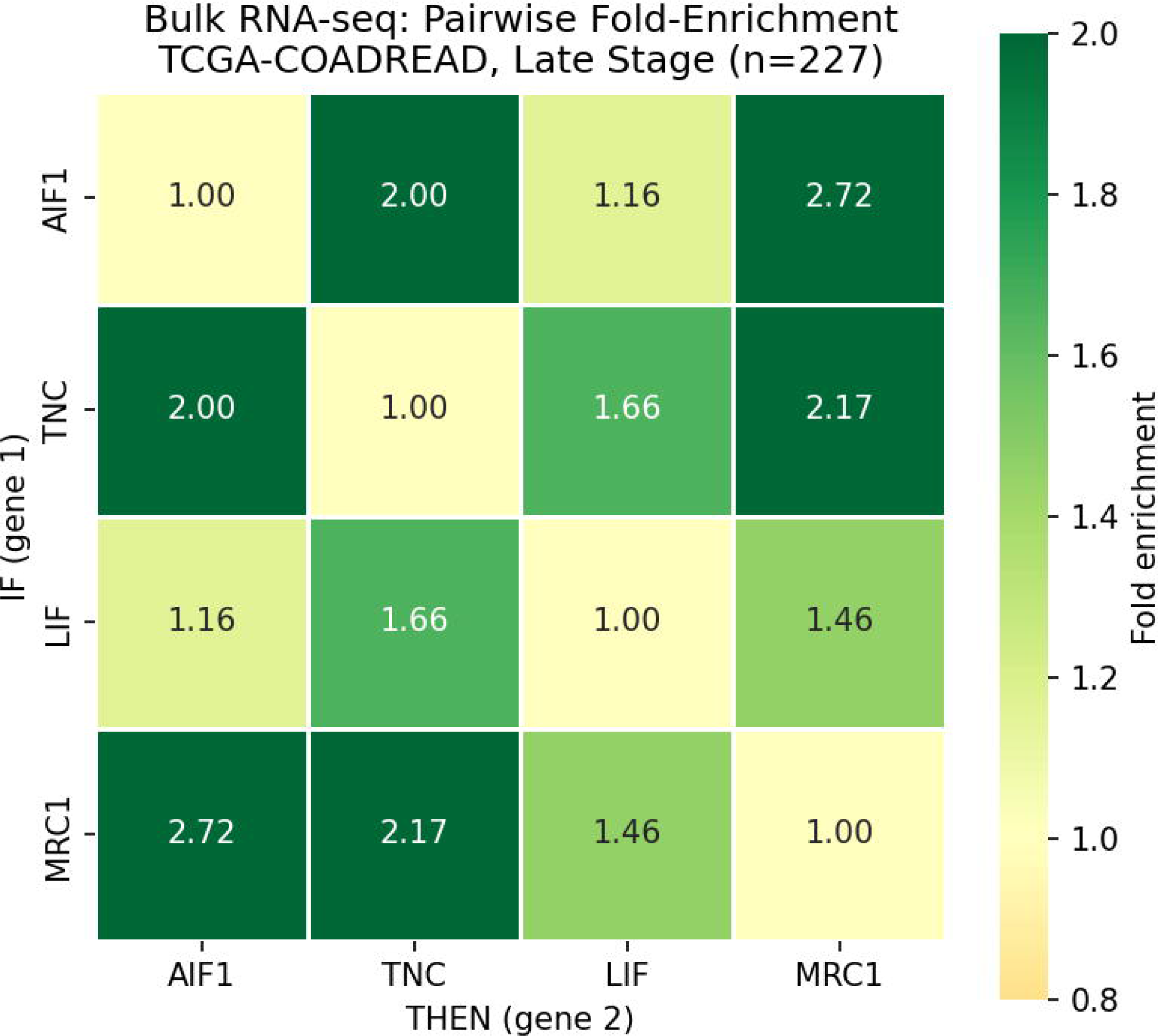
Pairwise Boolean co-occurrence fold-enrichment heatmap in bulk RNA-seq (Late Stage).

### MAS and glutamine metabolism dysregulation

We next examined whether metabolic pathway genes associated with the malate-aspartate shuttle (MAS) and glutamine metabolism show stage-dependent expression changes in TCGA-COADREAD. Genes included *GOT1, MDH1, GOT2*, and *MDH2* (MAS pathway) and *GLS*, *GLUD1*, and *GLUL* (glutamine pathway). Those gene expressions were compared between early-stage (n = 320) and late-stage (n = 227) samples using the Mann–Whitney U test, with Benjamini-Hochberg FDR correction across all seven genes (Table 4, Figure 2).

**Table 4.** Differential expression of MAS and glutamine pathway genes by tumor stage.

| Gene | Pathway | Early_median | Late_median | Delta | Direction | p_value | FDR |
| --- | --- | --- | --- | --- | --- | --- | --- |
| GOT1 | MAS | 1326.34 | 1243.6 | -82.74 | Downregulated | 0.00284678 | 0.009964 |
| MDH1 | MAS | 2618.66 | 2378.26 | -240.4 | Downregulated | 0.00066982 | 0.004689 |
| GOT2 | MAS | 3651.81 | 3511.97 | -139.8 | Downregulated | 0.142732551 | 0.199826 |
| MDH2 | MAS | 7056.54 | 6846.15 | -210.4 | Downregulated | 0.957745142 | 0.957745 |
| GLS | Glutamine | 2061.61 | 2141.86 | 80.25 | Upregulated | 0.27060509 | 0.315706 |
| GLUD1 | Glutamine | 3300.71 | 3130.32 | -170.39 | Downregulated | 0.024257294 | 0.0566 |
| GLUL | Glutamine | 6413.02 | 5982.65 | -430.37 | Downregulated | 0.124276817 | 0.199826 |

**Figure 2.**
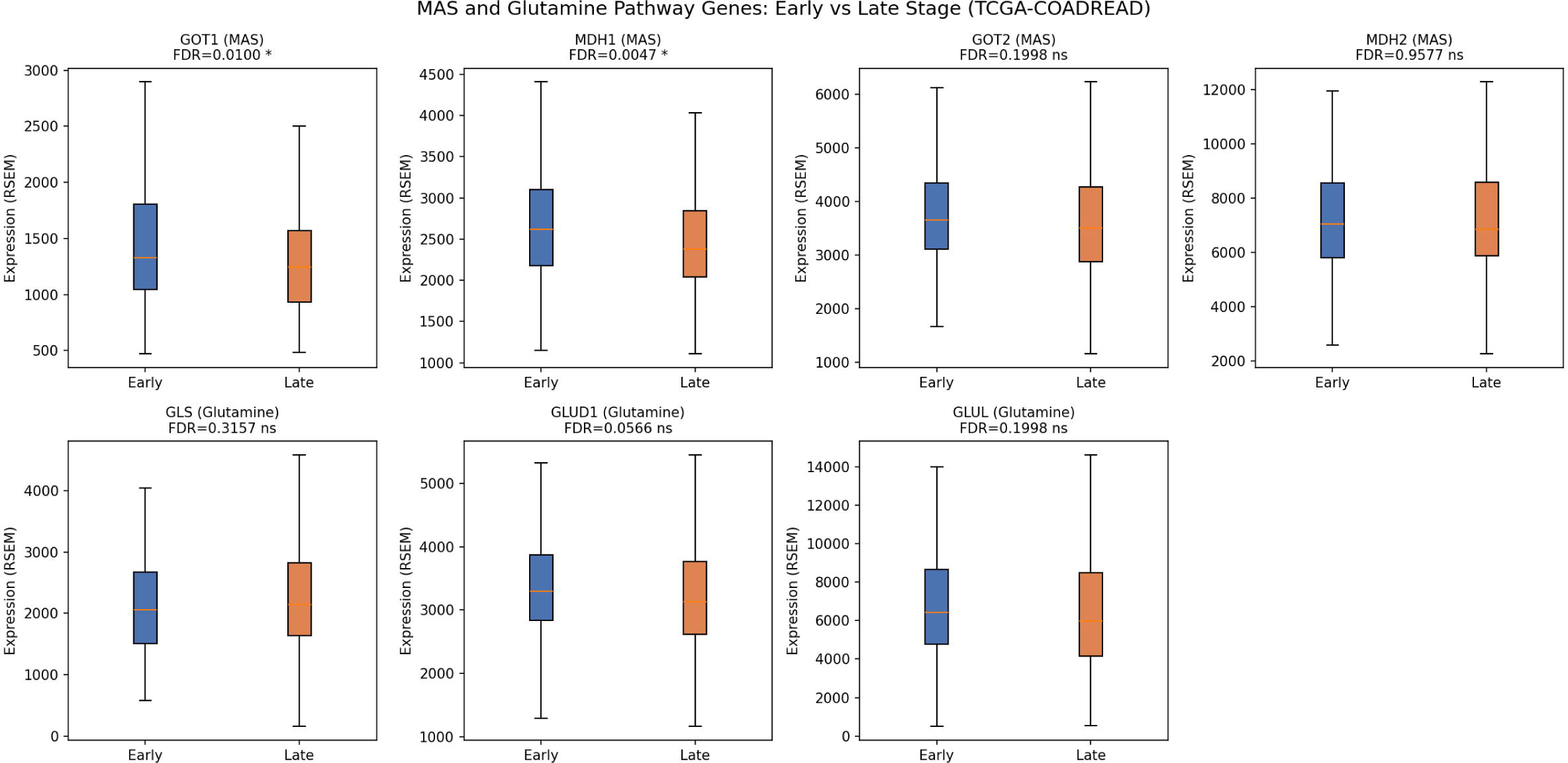
MAS and glutamine pathway gene expression across disease stages.

Only two MAS genes showed statistically significant downregulation in late-stage colorectal cancer. *GOT1* (Early median: 1,326 RSEM, Late: 1,244 RSEM, Delta =−83; FDR=0.010) and *MDH1* (Early: 2,619 RSEM, Late: 2,378 RSEM, Delta =−240; FDR=0.005). The remaining five genes did not show statistical differences after FDR correction (all FDR > 0.05), although *GLUD1* showed borderline downregulation (FDR = 0.057).

These observations that *GOT1* and *MDH1* were downregulated are consistent with the study by Ghirotto et al. (2026),^14^ which also pointed to reduced expression of these MAS genes in inflamed human gut tissue. Although the disease contexts differ, the shared downregulation of GOT1 and MDH1 suggests that alterations in the malate-aspartate shuttle (MAS) may occur across multiple pathological conditions in the gut. Further studies are needed to determine whether these changes reflect a common metabolic response or distinct diseases.

### Bulk survival analysis

To assess the clinical relevance of the selected four genes, we calculated a Boolean risk score for each patient by counting the number of genes with HIGH expression (top 25% of expression levels). The score ranged from 0 to 4. Higher scores indicated that more genes were highly expressed. Survival analysis included 545 patients with survival data, of whom 108 experienced an event (deaths). Patients with HIGH expression of all four genes (score = 4; n = 22) were classified as the High-Risk group, while all remaining patients (n = 523) were classified as the Low-Risk group.

Between the High-Risk and Low-Risk groups, Survival did not differ significantly in the Kaplan–Meier analysis (log-rank p = 0.103). In the Cox proportional hazards model, treating the risk score as a continuous variable, a higher risk score was associated with poorer overall survival (HR = 1.21 per unit increase, 95% CI: 1.03–1.41, p = 0.021; Supplementary Figure S2). Perhaps, limited statistical power in the small High-Risk group (n = 22) and only six events.

Overall, the four tumor budding genes showed only a weak association with patient survival and a marginally detectable prognostic signal in the bulk RNA-seq data; this marginal signal had fold-enrichments of 1.16–2.72x. The observed associations are weaker than the single-cell co-expression associations reported in the Single-Cell Validation Section.

### Single-Cell Validation

We used a publicly available scRNA-seq dataset from Pelka et al. (GSE178341) to examine whether the TB-module co-expression pattern is detectable at the single-cell level.^12^ This analysis included 26,057 cells and focused on histologically aggressive CRC subtypes (Mucinous and Medullary adenocarcinoma). These subtypes were selected as a proxy for aggressive TME biology, as direct tumor-budding annotation is not available in our selected dataset.

For each of the four TB-module genes, we defined a detection-based binary classification (count > 0). Detection rates were: *AIF1* was detected in 0.24% (62/26,057) of cells, *TNC* in 2.27% (591/26,057), *LIF* in 0.16% (41/26,057), and *MRC1* in 0.90% (235/26,057) of cells.

To evaluate Pairwise co-detection, we used Fisher’s exact tests for all 12 ordered gene pairs with Benjamini-Hochberg FDR correction (Table 5). Every pair showed statistically significant co-occurrence (all FDRs < 4×10⁻³⁹), with fold enrichments ranging from 31.0x to 184.5x. Notably, the strongest single-cell signal (*AIF1*–*LIF* at 184.5x) was not significant in bulk RNA-seq data (FE = 1.16x, FDR = 0.31), demonstrating that single-cell analysis can uncover critical biological signals that remain completely undetected in bulk analysis.

**Table 5.**
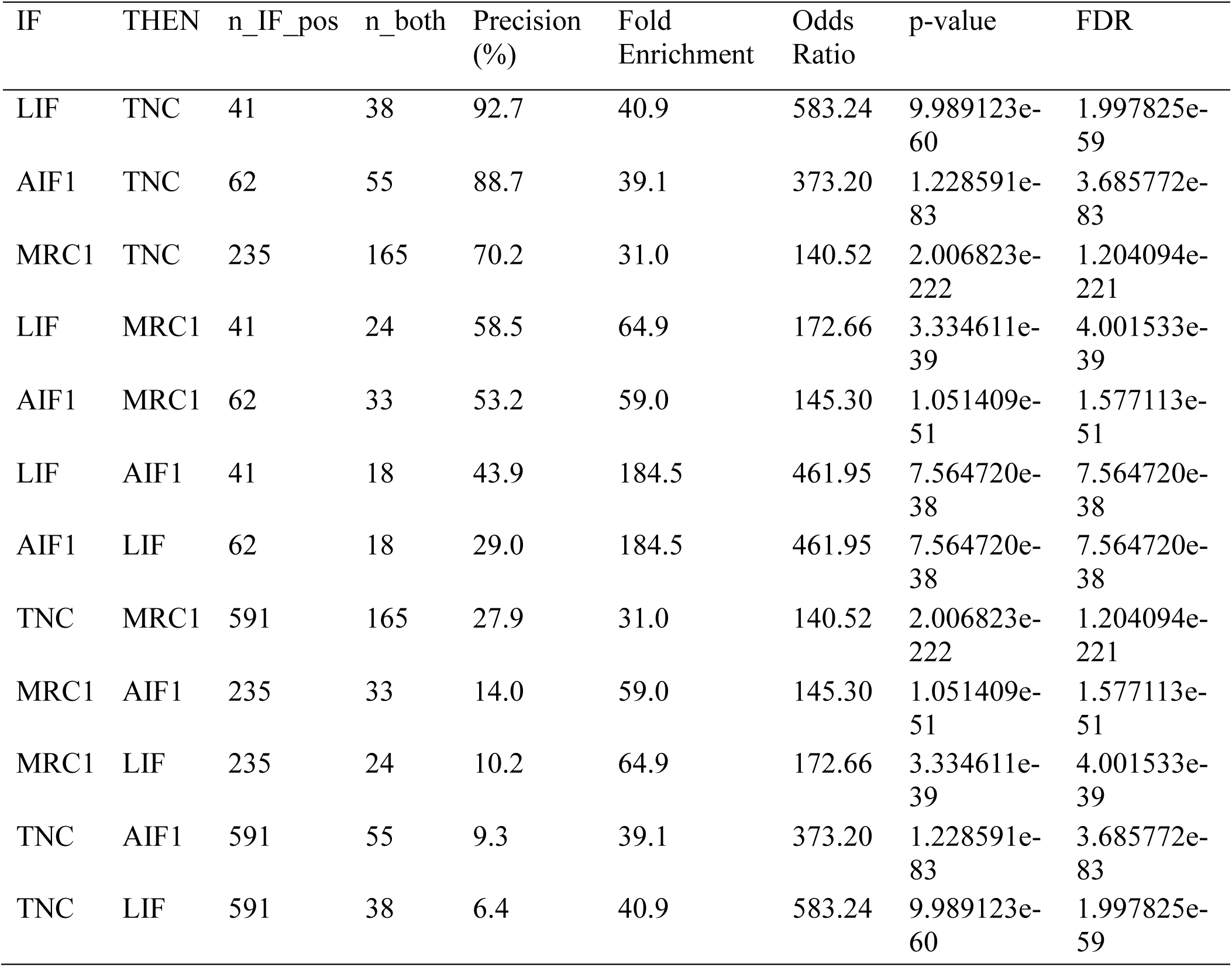
Single Cell Pairwise detection.

| IF | THEN | n_IF_pos | n_both | Precision (%) | Fold Enrichment | Odds Ratio | p-value | FDR |
| --- | --- | --- | --- | --- | --- | --- | --- | --- |
| LIF | TNC | 41 | 38 | 92.7 | 40.9 | 583.24 | 9.989123e-60 | 1.997825e-59 |
| AIF1 | TNC | 62 | 55 | 88.7 | 39.1 | 373.20 | 1.228591e-83 | 3.685772e-83 |
| MRC1 | TNC | 235 | 165 | 70.2 | 31.0 | 140.52 | 2.006823e-222 | 1.204094e-221 |
| LIF | MRC1 | 41 | 24 | 58.5 | 64.9 | 172.66 | 3.334611e-39 | 4.001533e-39 |
| AIF1 | MRC1 | 62 | 33 | 53.2 | 59.0 | 145.30 | 1.051409e-51 | 1.577113e-51 |
| LIF | AIF1 | 41 | 18 | 43.9 | 184.5 | 461.95 | 7.564720e-38 | 7.564720e-38 |
| AIF1 | LIF | 62 | 18 | 29.0 | 184.5 | 461.95 | 7.564720e-38 | 7.564720e-38 |
| TNC | MRC1 | 591 | 165 | 27.9 | 31.0 | 140.52 | 2.006823e-222 | 1.204094e-221 |
| MRC1 | AIF1 | 235 | 33 | 14.0 | 59.0 | 145.30 | 1.051409e-51 | 1.577113e-51 |
| MRC1 | LIF | 235 | 24 | 10.2 | 64.9 | 172.66 | 3.334611e-39 | 4.001533e-39 |
| TNC | AIF1 | 591 | 55 | 9.3 | 39.1 | 373.20 | 1.228591e-83 | 3.685772e-83 |
| TNC | LIF | 591 | 38 | 6.4 | 40.9 | 583.24 | 9.989123e-60 | 1.997825e-59 |

We used Mann-Whitney U tests to exclude technical doublet artifacts, which can artificially increase co-detection rates by capturing two cells. We compared UMI counts between gene-positive and gene-negative cells for each of the four genes. We did not find any significant increase in total UMI counts in gene-positive cells across all four cases (all Mann-Whitney p > 0.50; Figure 3). So these are not doublets.

**Figure 3.**
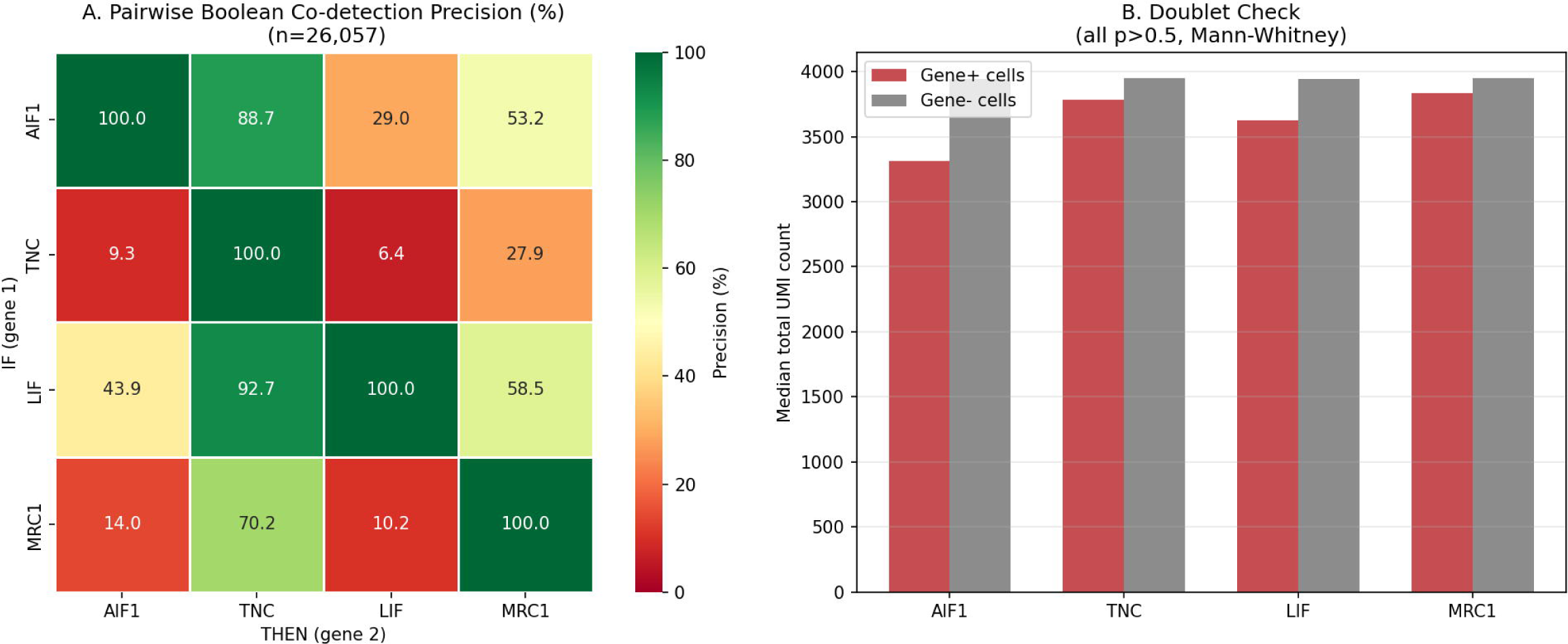
Single-cell Boolean co-detection analysis in aggressive-subtype CRC. Two-panel figure. **(a)** Heatmap of pairwise Boolean co-detection precision. (b) Doublet artifact check. Grouped bar chart showing median total UMI count for gene-positive (red) versus gene-negative (gray) cells for each of the four genes.

To assess whether the pooled co-expression patterns were influenced by individual donor differences, we analyzed gene detection separately within each patient. All four TB-module genes were detected only in two of the eight patients (C106 and C110). Importantly, the strong pairwise gene enrichment remained above baseline when these two patients were analyzed separately. For example, *AIF1*–*LIF* fold enrichment was 20.9-fold in C106 and 45.4-fold in C110, while *AIF1*–*TNC* enrichment was 6.6- and 6.7-fold, respectively. Across all six unique gene pairs, within-patient fold enrichment ranged from 4.6 to 20.9 in C106 and from 6.0 to 45.4 in C110 (Supplementary Table 3). These results demonstrate that the observed co-detection pattern was present within both patients and was not solely a consequence of pooling cells across donors.

### Bulk vs single-cell comparison

A comparison of the bulk and single-cell results demonstrated that pairwise co-expression was much stronger in the single-cell data (Table 6, Figure 4). Overall, these results show that the TB-module co-expression pattern was more clearly detected in the single-cell data.

**Table 6.** Bulk vs Single-cell comparison table.

| Pair | Bulk FE (FDR) | Single-cell FE (FDR) | SC/Bulk ratio |
| --- | --- | --- | --- |
| AIF1-LIF | 1.16x (ns, 0.31) | 184.5x (7.6e-38) | ~159x |
| LIF-MRC1 | 1.46x (0.037) | 64.9x (4.0e-39) | ~44x |
| LIF-TNC | 1.66x (0.0028) | 40.9x (2.0e-59) | ~25x |
| AIF1-TNC | 2.00x (1.2e-5) | 39.1x (3.7e-83) | ~20x |
| MRC1-TNC | 2.17x (5.9e-8) | 31.0x (1.2e-221) | ~14x |
| AIF1-MRC1 | 2.72x (1.7e-12) | 59.0x (1.6e-51) | ~22x |

**Figure 4.**
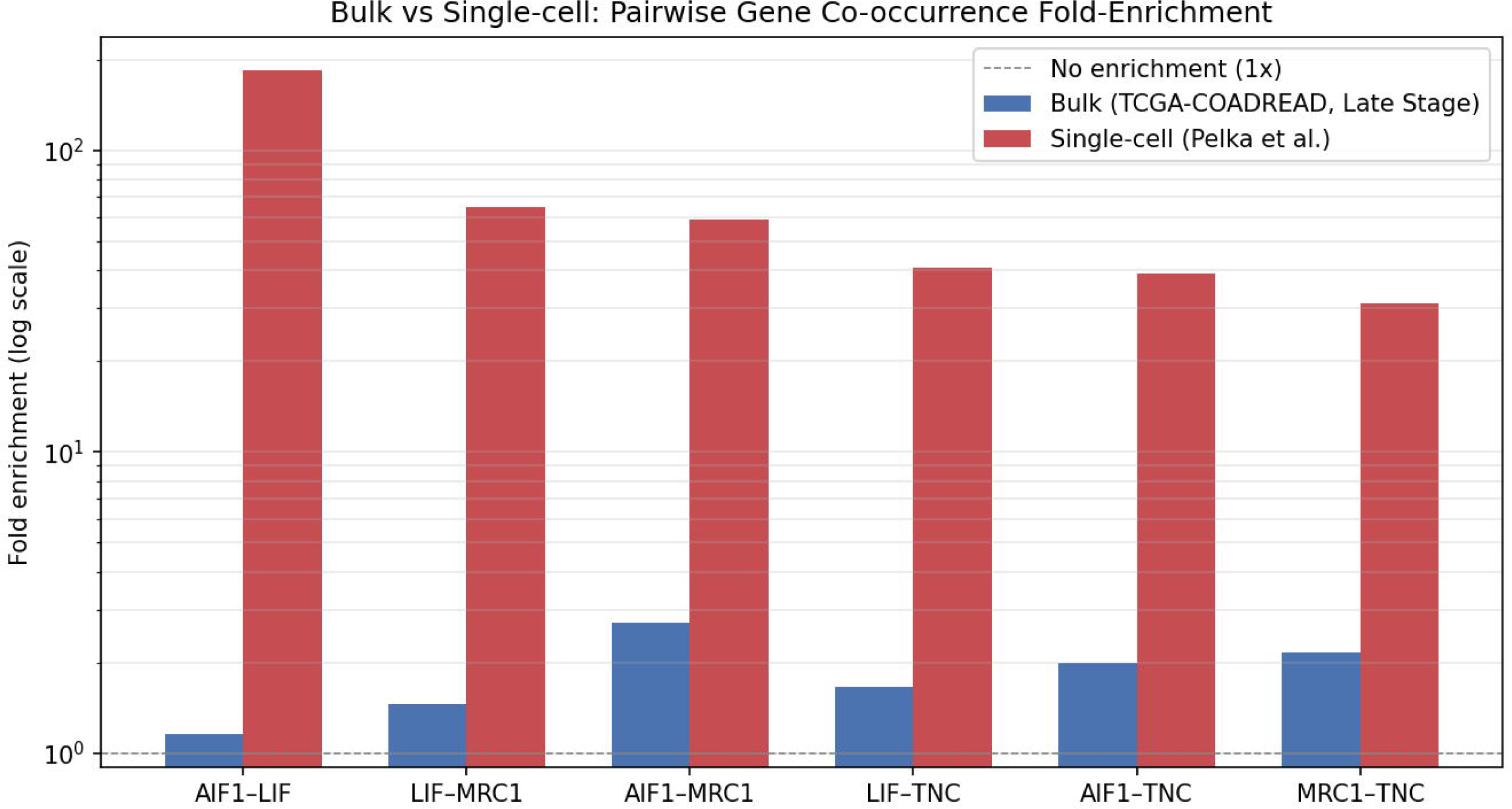
Comparison of pairwise fold-enrichment between bulk RNA-seq and single-cell RNA-seq.

## Discussion

This study examined whether a four-gene tumor-budding (TB)-associated module consisting of *AIF1, LIF, MRC1*, and *TNC* could also be detected as a co-expression pattern in bulk RNA-seq data. These genes were previously identified by Phoenix virtual spatial transcriptomics as enriched in tumor-budding regions of CRC.^8^ Our findings demonstrate that the co-expression pattern was much weaker in bulk RNA-seq than in single-cell data. That being said, averaging gene expression across many cell types in bulk tissue may mask cell-specific co-expression signals.^15^

Across all six gene pairs, fold enrichments exceeded bulk fold enrichments by 14x to 159x. Most specifically, *AIF1*–*LIF* pairs showed the strongest single-cell association (FE = 184.5, FDR ≈ 10⁻³⁸) but no statistically significant co-occurrence in bulk RNA-seq (FE = 1.16, FDR = 0.31). This bulk invisibility is not a consequence of insufficient sample size (TCGA-COADREAD provides n=547 patients), with n=26,057 cells in the single-cell dataset. That may reflect differences in biological information between the two approaches, because bulk RNA-seq measures average gene expression across many cell types within a tumor sample, whereas single-cell RNA-seq measures expression in individual cells.^16–18^ This interpretation is also consistent with the low single-cell detection rates observed for the four genes (0.16–2.27%), which are vastly outnumbered by signal-negative cells, their co-expression relationships are diluted beyond the threshold of bulk detectability.

At the single-cell level, the four tumor budding genes showed tightly connected co-expression and significant pairwise associations in the aggressive-subtype CRC cells (all FDR < 4 × 10⁻³⁹). *AIF1* and *MRC1* are associated with macrophage populations, and *LIF* is a cytokine frequently produced by tumor-associated macrophages and cancer-associated fibroblasts. *TNC* is an extracellular matrix glycoprotein commonly found in tumor stroma and the invasive front. *TNC* acts as a co-expression hub to which all three myeloid genes converge (precision 70–93%). Together, these findings suggest that the observed co-expression may reflect interactions among myeloid/ macrophage and stromal components of the tumor microenvironment.^19–21^

Notably, we excluded technical doublet artifacts. Gene-positive cells showed no elevation in total UMI count relative to gene-negative cells for any of the four genes (Mann-Whitney p > 0.50 for all).

We used UniversalEPI to analyze gene expression in HCT116 cells, a standard epithelial colon cancer cell line.^11^ This test gave us clear physical proof of how different genes work in specific cell types. *TNC* and *LIF* showed significant predicted interactions (n = 61 and 33, respectively), with maximum Z-scores up to 5.8. This means these genes are actively working and turned on. In contrast, the switch for the *AIF1* gene was missing from the HCT116 map, meaning no control links were found at this location. This absence is expected because *AIF1* is a specialized gene expressed only in immune cells, such as macrophages.^22,23^ In colon cancer cells, this gene’s region is closed, meaning it can only be opened inside an immune cell environment. Since HCT116 is an epithelial colorectal cancer cell line, this finding points toward a cell-type-specific setup for *AIF1*. However, additional chromatin accessibility and cell-type-specific EPI data would be required to confirm this interpretation. Ultimately, this discovery provides a clear explanation for a common problem in cancer testing: when we test a whole tumor sample (bulk tissue) and find *AIF1*, it does not mean the cancer cells themselves are changing. Instead, it serves as a clear biological signal that immune cells have entered the tumor. It also demonstrates why it is so difficult to catch *AIF1* working together with cancer genes in bulk tissue tests.

Beyond the TB-module analysis, we found lower expression of two malate-aspartate shuttle (MAS) genes, *GOT1* (FDR = 0.010) and *MDH1* (FDR = 0.005), in late-stage compared with early-stage colorectal cancer (CRC). The MAS pathway acts like a cellular transport system, moving vital energy molecules across the mitochondrial membrane to keep a healthy NAD⁺/NADH balance and support normal mitochondrial energy production (OXPHOS).^24,25^ When these two enzymes are downregulated in CRC, it is associated with a major metabolic shift. It suggests that as the tumor progresses, it relies less on mitochondrial respiration and shifts its energy production toward high-glucose consumption. This classic cancer hallmark is known as the Warburg effect.^26,27^ Recently identified MAS suppression, specifically involving *GOT1* and *MDH1*.^14^ This paper showed the exact same pattern: shutting down *GOT1* and *MDH1* is a core signature of inflamed gut tissues, seen in both Parkinson’s disease models and ulcerative colitis biopsies. Although the disease contexts differ, this suggests that *GOT1* and *MDH1* represent a shared metabolic weak spot when gut tissues face severe cellular stress. While these data provide strong clues, Further studies using metabolomics and functional experiments are needed.

However, we acknowledged several limitations. First, the single-cell analysis used aggressive types of colon cancer (mucinous and medullary adenocarcinoma) as a proxy because direct tumor-budding annotations were not available in the dataset.^12^ Our findings should be viewed as supporting, rather than directly validating, the original Phoenix TB-zone signature. Second, the Boolean threshold used in the single-cell analysis (count > 0) represents gene detection rather than high gene expression. This can be affected by technical factors like sequencing depth and gene-capture efficiency. Third, our predicted enhancer–promoter interaction analysis is based on just one cell line (HCT116). Therefore, these results are exploratory and descriptive rather than exact statistical proof. Fourth, the patient survival analysis showed only a weakly significant association (HR = 1.21, p = 0.021), and the high-risk group was small (n = 22). These survival findings require validation in a larger independent patient group. Another limitation is that all four TB-module genes were detected in only two out of the eight patients within the single-cell cohort. Although within-patient analysis showed consistent pairwise enrichment in both patients, the small number of patients still prevents assessment of how broadly this pattern is shared across CRC tumors. Therefore, future validation in larger independent single-cell datasets with appropriate cell-type annotations will be essential for confirmation.

Finally, all the findings presented here are based on publicly available datasets and computational approaches. While these models provide valuable insights, real laboratory experiments using patient-derived models remain an important next step to confirm our discoveries about the immune system and cellular metabolism.

## Conclusion

Our study demonstrates that the four-gene TB-associated module (*AIF1, LIF, MRC1*, and *TNC*) was clearly detected at the single-cell level compared to bulk RNA-seq data. Pairwise co-expression signals were stronger in single-cell data, while bulk analysis showed weaker associations and limited prognostic value. Importantly, while this TB-module co-expression was more clearly resolved at single-cell resolution, its detection varied substantially across individual patients. Additionally, predicted enhancer–promoter interaction analysis suggested that some of these genes may be regulated in a cell-type-dependent manner. The lower expression of the MAS genes *GOT1* and *MDH1* in late-stage CRC suggests that metabolic changes may accompany disease progression. Overall, our findings highlight the importance of choosing the right transcriptomic platform when studying gene expression patterns in specific cell types or tumor microenvironments. Further validation in independent cohorts and experimental models will be needed to confirm these findings and determine their biological and clinical relevance.

## Method

### Promoter Region Extraction

To identify the promoter regions of the eight candidate genes (*AIF1, TNC, LIF, MRC1, GOT1*, *MDH1, GOT2, MDH2*), we used the Ensembl GRCh38 genome annotation, downloaded from the FTP server (https://ftp.ensembl.org/pub/release-110/gtf/homo_sapiens/). For each gene, we identified the transcription start site (TSS) using gene-level GTF annotations. Promoter intervals were defined relative to the TSS, with a 2,500-bp window per gene. This window spanned 2,000 bp upstream to 500 bp downstream of the TSS for both forward- and reverse-strand genes.

### Predicted enhancer-promoter interaction analysis

The HCT116 colorectal cancer cell EPI (enhancer-promoter interaction) data were downloaded from the BoevaLab UniversalEPI server.^28^ We used the bigBed-format interaction file (hct116_z.bb).^29^ This file lists predicted pairs of enhancer-promoter interactions. Each record includes two anchor regions (5 kb bins) and an interaction Z-score representing the interaction strength. We mapped each promoter region to fixed 5-kb genomic bins (bin size = 5,000 bp). We searched a ±2 Mb window around each promoter to extract all matching chromatin interaction entries from the bigBed file using the pyBigWig library (v0.3.18). An interaction was linked to a gene if either of its two anchors matched the gene’s 5 kb promoter bins. If their Z-score was above 1.645 (p < 0.05), we considered it significant. For each gene, we counted the total number of interactions, the maximum Z-score, the average Z-score, and the number of significant interactions.

### TCGA-COADREAD bulk RNA-seq processing and Boolean co-occurrence analysis

Bulk RNA-seq data (RSEM-normalized) and matched clinical data were obtained from cBioPortal (https://www.cbioportal.org). Stages were binarized based on Early (Stage I–II) or Late (Stage III–IV). A total of 547 patients were retained (Early: 320; Late: 227) after stage filtering.

Gene expression levels were binarized using the 75th percentile of sample-wide expression as the threshold. All 547 patients were computed regardless of stage, such that patients in the top quartile of expression were classified as HIGH (bool = 1) and all others as LOW (bool = 0). This threshold was selected over the commonly used median split to avoid artificially forcing a 50% activation rate, thereby preserving biological meaningfulness. The specific percentile thresholds were: *AIF1* = 253.88, *TNC* = 2,984.37, *LIF* = 1,043.78, and *MRC1* = 332.55.

• **Early Stage (n = 320):** *AIF1*: 27.2%; *TNC*: 24.4%; *LIF*: 26.9%; *MRC1*: 25.6%.

• **Late Stage (n = 227):** *AIF1*: 22.0%; *TNC*: 26.0%; *LIF*: 22.5%; *MRC1*: 24.2%.

Pairwise co-occurrence was assessed in the Late-stage patients (n = 227). A total of 12 directional gene pairs were tested from the four selected genes (*AIF1, TNC, LIF*, and *MRC1*). For each ordered pair, a one-sided Fisher’s exact test was performed. To calculate the fold enrichment, we compared how often Gene 2 was highly expressed in patients who already had high levels of Gene 1 against the overall baseline rate of Gene 2 in all Late-stage patients. To control the false discovery rate (FDR) from multiple comparisons, we used the Benjamini-Hochberg (BH) procedure. All statistical analyses were implemented in Python and the statsmodels library.

### MAS and glutamine pathway differential expression

To compare the expression levels of seven metabolic genes (*GOT1, MDH1, GOT2, MDH2* known for malate-aspartate shuttle; *GLS, GLUD1, GLUL* known for glutamine metabolism) between Early-stage and Late-stage patients,^14,30^ Two-sided Mann-Whitney U tests were used for each gene. We measured the change in expression by calculating the difference between the medians of the two groups (Late-stage and Early-stage). Multiple testing correction was applied using the Benjamini-Hochberg (BH) procedure across all tests. Genes with FDR < 0.05 were considered significant.

### Survival analysis

Survival analysis was performed using the lifelines library (v0.27). Survival time was taken from the OS_MONTHS field in the TCGA-COADREAD clinical data file. Event status was derived from OS_STATUS (“1:DECEASED” = event, “0:LIVING” = censored). Data with missing OS_MONTHS or OS_STATUS were excluded; therefore, 545 patients with valid survival data (108 deaths, 437 Alive).

A four-gene Boolean risk score was created by summing the Boolean HIGH indicators for *AIF1, TNC, LIF,* and *MRC1* for each patient (score range 0–4).

1. **High-Risk Group (n = 22):** Patients who had all four genes highly expressed at the same time (score = 4).
2. **Low-Risk Group (n = 523):** All other patients (score between 0 and 3).

We used a continuous Cox proportional hazards regression model to evaluate the risk score (0 to 4) as a single predictor of survival. This model calculated the hazard ratio (HR), 95% confidence intervals, and p-values without adding any other clinical variables. All survival computations were carried out using Python’s lifelines library.

### Single-cell RNA-seq processing and co-occurrence analysis

Single-cell RNA-sequencing data were downloaded from a published dataset (Pelka et al., 2021) via the NCBI GEO database as a CellRanger-format HDF5 file and matched metadata.^12,31^ Cells were filtered to retain only aggressive subtypes (Mucinous adenocarcinoma and Medullary adenocarcinoma); that filtering left us a total of 26,057 cells. Raw count data for the four genes (*AIF1, TNC, LIF, MRC1*) were extracted from the HDF5 sparse matrix file (43,113 genes × 370,115 cells) using h5py (v3.9), gene indices identified by exact name matching against the features/name array.

Single-cell data often has dropouts, where low-expressing genes accidentally show up as zero. We used a simple detection-based rule to classify expression:

1. **High Group (1):** raw count greater than zero.
2. **Low Group (0):** Any cell with a count of exactly zero.

By following this rule, the baseline cell detection rates were: *AIF1* (0.24%), *TNC* (2.27%), *LIF* (0.16%), and *MRC1* (0.90%).

Pairwise co-detection was assessed using Fisher’s exact tests for all 12 gene pairs, with BH-FDR correction. To determine fold-enrichment, we calculated the co-detection rate of Gene 2 within Gene 1-High cells and divided it by the baseline detection rate of Gene 2 across all 26,057 cells.

Next, we ran a safety check to make sure that our results were biologically sound and not a technical error called “cell doublets” (When two different cells accidentally get trapped inside the same oil droplet during lab processing). Doublets are actually a combination of two cells, they naturally show a higher total gene count (total UMI count) than a single cell.^32,33^ We used a Mann-Whitney U test to compare the UMI counts between gene-positive and gene-negative cells. The lack of gene inflation in our positive cells shows that our data is free from doublets. All analyses were performed using Python.

## Data Availability

Single-Cell RNA-sequencing data were obtained from the NCBI Gene Expression Omnibus (GEO) database under accession number GSE178341. Bulk RNA-seq and Clinical Profiles and matched survival outcomes were retrieved directly via the cBioPortal from the Cancer Genomics platform (cbioportal.org). The genomic annotations, Transcription Start Sites (TSS), and 2,500 bp flanking promoter sequences used for structural mapping were extracted using the official Ensembl GRCh38 reference system.

## Code Availability

The scripts used during the current study are available from the corresponding author upon reasonable request.

## Author Contributions

Pritom Sarkar and Prapthi Sakar conceptualized the idea. Pritom Sarkar designed the multi-omics workflow, developed the transcriptomic and statistical software architecture, performed the formal data analysis, and generated the figures and illustrations. He was also responsible for investigating and drafting the original manuscript. Prapthi Sarkar conducted the data curation, implemented the biological annotation mapping across clinical references, performed functional pathway validations for the target metabolic signatures, and contributed significantly to writing, reviewing, and editing the final manuscript.

## Corresponding Author

Correspondence to Pritom Sarkar.

## Additional Information

### Supplementary Information

Supplementary material accompanies this paper online.

