## Supplementary Information PDF for "Cross-Platform Evaluation of a Tumor-Budding-Associated Gene Signature in Colorectal Cancer"

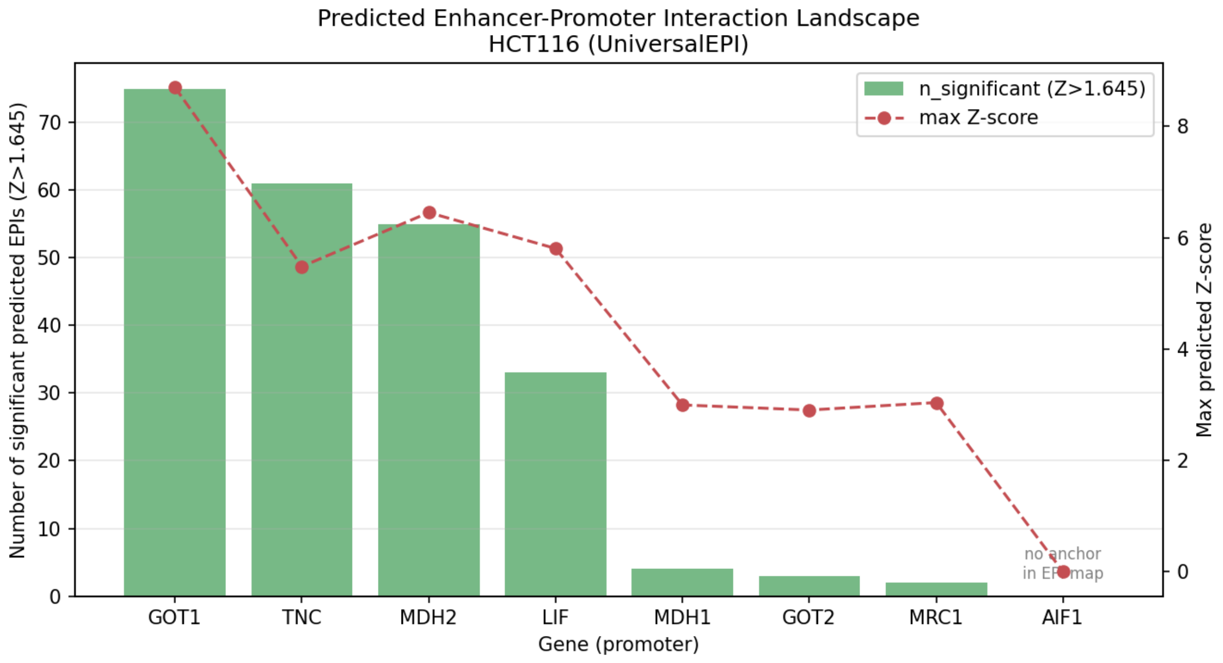


Fig. S1. Enhancer-promoter interaction (EPI) landscape at promoter regions of selected genes in HCT116 cells.

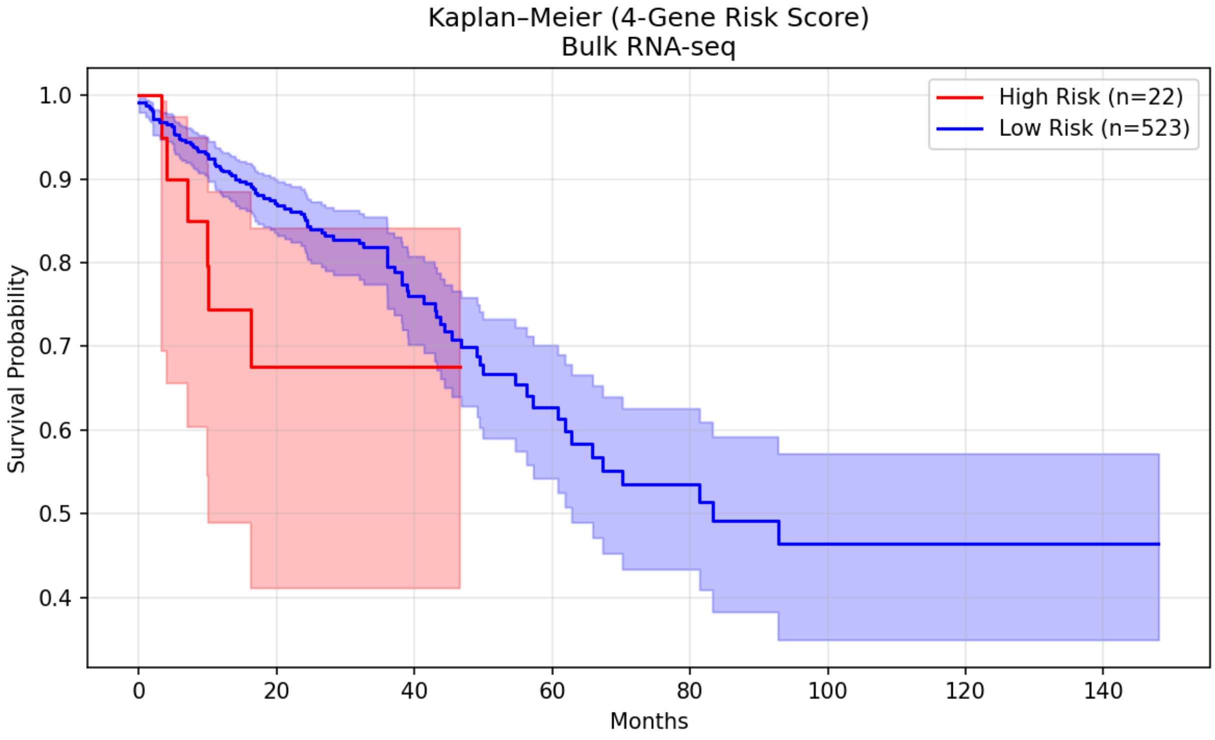


Fig. S2. Kaplan–Meier survival curves for 545 colorectal cancer patients stratified by the 4-gene Boolean risk score (High: n = 22; Low: n = 523). Log-rank p = 0.103. Cox PH with continuous risk score: HR = 1.21 (95% CI: 1.03–1.41), p = 0.021.

Supplementary Table 1: Promoter Region

| **Gene Symbol** | **Chromosome** | **Start Position** | **End Position** |
|---|---|---|---|
| AIF1 | 6 | 31,613,217 | 31,615,717 |
| GOT1 | 10 | 99,430,124 | 99,432,624 |
| GOT2 | 16 | 58,733,842 | 58,736,342 |
| LIF | 22 | 30,246,259 | 30,248,759 |
| MDH1 | 2 | 63,586,609 | 63,589,109 |
| MDH2 | 7 | 76,046,051 | 76,048,551 |
| MRC1 | 10 | 17,807,348 | 17,809,848 |
| TNC | 9 | 115,117,707 | 115,120,207 |

Supplementary Table 2: MAS and glutamine pathway genes between Early and Late stages.

| Gene | Pathway | Early_median | Late_median | Delta | Direction | p_value | FDR |
|---|---|---|---|---|---|---|---|
| GOT1 | MAS | 1326.34 | 1243.60 | -82.74 | ↓ in Late | 0.002847 | 0.009964 |
| MDH1 | MAS | 2618.66 | 2378.26 | -240.40 | ↓ in Late | 0.000670 | 0.004689 |
| GOT2 | MAS | 3651.81 | 3511.97 | -139.84 | ↓ in Late | 0.142733 | 0.199826 |
| MDH2 | MAS | 7056.54 | 6846.15 | -210.39 | ↓ in Late | 0.957745 | 0.957745 |
| GLS | Glutamine | 2061.61 | 2141.86 | 80.25 | ↑ in Late | 0.270605 | 0.315706 |
| GLUD1 | Glutamine | 3300.71 | 3130.32 | -170.39 | ↓ in Late | 0.024257 | 0.056600 |
| GLUL | Glutamine | 6413.02 | 5982.65 | -430.37 | ↓ in Late | 0.124277 | 0.199826 |

Supplementary Table 3: Per-patient Co-occurrence Analysis of in Colorectal Cancer Patients (C106 and C110)

| Gene Pair | C106 n_IF | C106 Precision (%) | C106 Baseline (%) | C106 FE | C110 n_IF | C110 Precision (%) | C110 Baseline (%) | C110 FE |
|---|---|---|---|---|---|---|---|---|
| AIF1→TNC | 28 | 89.3 | 13.4 | 6.6 | 34 | 88.2 | 13.1 | 6.7 |
| AIF1→LIF | 28 | 25.0 | 1.2 | 20.9 | 34 | 32.4 | 0.7 | 45.4 |
| AIF1→MRC1 | 28 | 60.7 | 6.0 | 10.1 | 34 | 47.1 | 4.7 | 10.0 |
| TNC→AIF1 | 259 | 9.7 | 1.5 | 6.6 | 332 | 9.0 | 1.3 | 6.7 |
| TNC→LIF | 259 | 8.1 | 1.2 | 6.8 | 332 | 5.1 | 0.7 | 7.2 |
| TNC→MRC1 | 259 | 27.4 | 6.0 | 4.6 | 332 | 28.3 | 4.7 | 6.0 |
| LIF→AIF1 | 23 | 30.4 | 1.5 | 20.9 | 18 | 61.1 | 1.3 | 45.4 |
| LIF→TNC | 23 | 91.3 | 13.4 | 6.8 | 18 | 94.4 | 13.1 | 7.2 |
| LIF→MRC1 | 23 | 56.5 | 6.0 | 9.4 | 18 | 61.1 | 4.7 | 13.0 |

| | | | | | | | | |
|---|---|---|---|---|---|---|---|---|
| MRC1→AIF1 | 116 | 14.7 | 1.5 | 10.1 | 119 | 13.4 | 1.3 | 10.0 |
| MRC1→TNC | 116 | 61.2 | 13.4 | 4.6 | 119 | 79.0 | 13.1 | 6.0 |
| MRC1→LIF | 116 | 11.2 | 1.2 | 9.4 | 119 | 9.2 | 0.7 | 13.0 |

*Note: n_IF = number of cells expressing the first gene; Precision = percentage of n_IF cells that also express the second gene; Baseline = overall expression frequency of the second gene; FE = fold enrichment (Precision / Baseline). Patient C106: n = 1926 cells; Patient C110: n = 2527 cells.*
